# Binocular combination in the autonomic nervous system

**DOI:** 10.1101/2024.06.04.597314

**Authors:** Federico G. Segala, Aurelio Bruno, Joel T. Martin, Anisa Y. Morsi, Alex R. Wade, Daniel H. Baker

## Abstract

Pupil diameters are regulated by the autonomic nervous system, which combines light signals across the eyes independently of the visual cortex. Distinct classes of retinal photoreceptor are involved in this process, with cones and rods driving the initial constriction and intrinsically photosensitive retinal ganglion cells maintaining diameter over prolonged time periods. We investigated binocular combination by targeting different photoreceptor pathways using a novel binocular multiprimary system to modulate the input spectra via silent substitution. At the first harmonic of the modulation frequency, luminance and S-cone responses showed strong binocular facilitation, and weak interocular suppression. Melanopsin responses were invariant to the number of eyes stimulated. The L-M pathway involved binocular inhibition, whereby responses to binocular stimulation were weaker than for monocular stimulation. The second harmonic involved strong interocular suppression in all pathways, but with some evidence of binocular facilitation. Our results are consistent with a computational model of binocular signal combination (implemented in a Bayesian hierarchical framework), in which the weight of interocular suppression differs across pathways. We also find pathway differences in response phase, consistent with different lag times for phototransduction. These results provide evidence that binocular interactions in the pupillary pathway differ across photoreceptor-directed modulations and across harmonic components of the response.

## Introduction

The autonomic nervous system regulates many involuntary bodily processes, including the constriction and dilation of the pupils in response to light [1]. The anatomical pathway from the retina to the subcortical nuclei controlling the pupillary light response (PLR) is well established: it includes the olivary pretectal nucleus (OPN), the Superior Cervical ganglion and the Edinger-Westphal nucleus, which project to the iris sphincter muscles that directly control the pupil size [1]. Evidence of a binocular component to the PLR is shown by the consensual response of the pupil (stimulation of one eye will cause constriction of the other eye) [2]. The anatomical segregation of the subcortical pathway from the rest of the brain means that this binocular combination of signals must occur independently of the cortical processes of binocular integration required for visual perception. Our recent work [3] has shown that the algorithm underlying binocular combination of light in the pupil pathway differs from that in the cortex. Here we extend this paradigm to compare binocular combination across signals from different photoreceptor classes that feed into the autonomic nervous system.

Different classes of retinal photoreceptors, including cones, rods and melanopsin-containing intrinsically photosensitive retinal ganglion cells (ipRGCs), are directly involved in controlling and maintaining the size of the pupils [4–8]. Cones drive the initial rapid constriction of the pupils [9], while the slower and longer activation of the ipRGCs maintains constriction over a prolonged period of time and regulates the postillumination pupillary response [4, 10]. The ipRGCs are a recently discovered photoreceptor class [11] that express the photopigment melanopsin, and are involved in the regulation of the circadian rhythm [12, 13], forming a major input to the OPN [14]. The first direct evidence of the involvement of the ipRGCs in the PLR was shown in melanopsin knockout mice [15], resulting in the loss of the intrinsic photosensitivity of the cells and a reduced pupil constriction. Similar behaviour was later observed in primates and humans [16] using silent substitution (see Methods), where it was shown that the PLR continues during light presentation even when cone and rod signalling is blocked, indicating the primary role of the ipRGCs in maintaining pupil constriction over a prolonged time.

Binocular combination has been extensively studied in visual perception, where it is mediated by neurons in primary visual cortex [17]. For pattern vision, binocular summation occurs at threshold, such that a lower contrast is required to detect a target shown to both eyes than a target shown to one eye [18, 19]. At higher contrasts, the phenomenon of ‘ocularity invariance’ is observed, in which the response to monocularly- and binocularly-presented patterns is equal [20, 21]. This is explained by a process of interocular suppression that cancels out the additional excitatory drive caused by stimulating two eyes. Our recent work [3] showed that cortical signal combination for luminance flicker is substantially more linear than for spatial patterns, whereas there is evidence of interocular suppression in the pupil pathway. Here we measure the amplitude of pupil modulations in response to flickering stimuli presented as light flux, or directed towards the L-M cones, S-cones and melanopsin (see Figure 1e-l). Our design uses three comparisons, each estimating a distinct quantity. To obtain an objective measure of pupil responses, we presented the two eyes with slightly different temporal frequencies (0.5 and 0.4 Hz). Responses at these two frequencies fall in separate bins of the Fourier spectrum and so this ‘frequency tagging’ lets us recover the response driven by one eye while the other eye supplies a concurrent, separable signal. Comparing monocular and binocular stimulation measures the net effect of adding a second eye, but cannot by itself separate additional excitation from interocular suppression. Therefore we used a dichoptic mask, in which the target eye is stimulated while the other eye views a fixed-contrast modulation. This allows us to isolate the suppression that one eye exerts on the response to the other. Together these comparisons allow us to test whether binocular facilitation and interocular suppression differ across photoreceptor-directed modulations. We interpret the results using a contemporary model of binocular vision [22], fitted within a hierarchical Bayesian framework.

**Figure 1:**
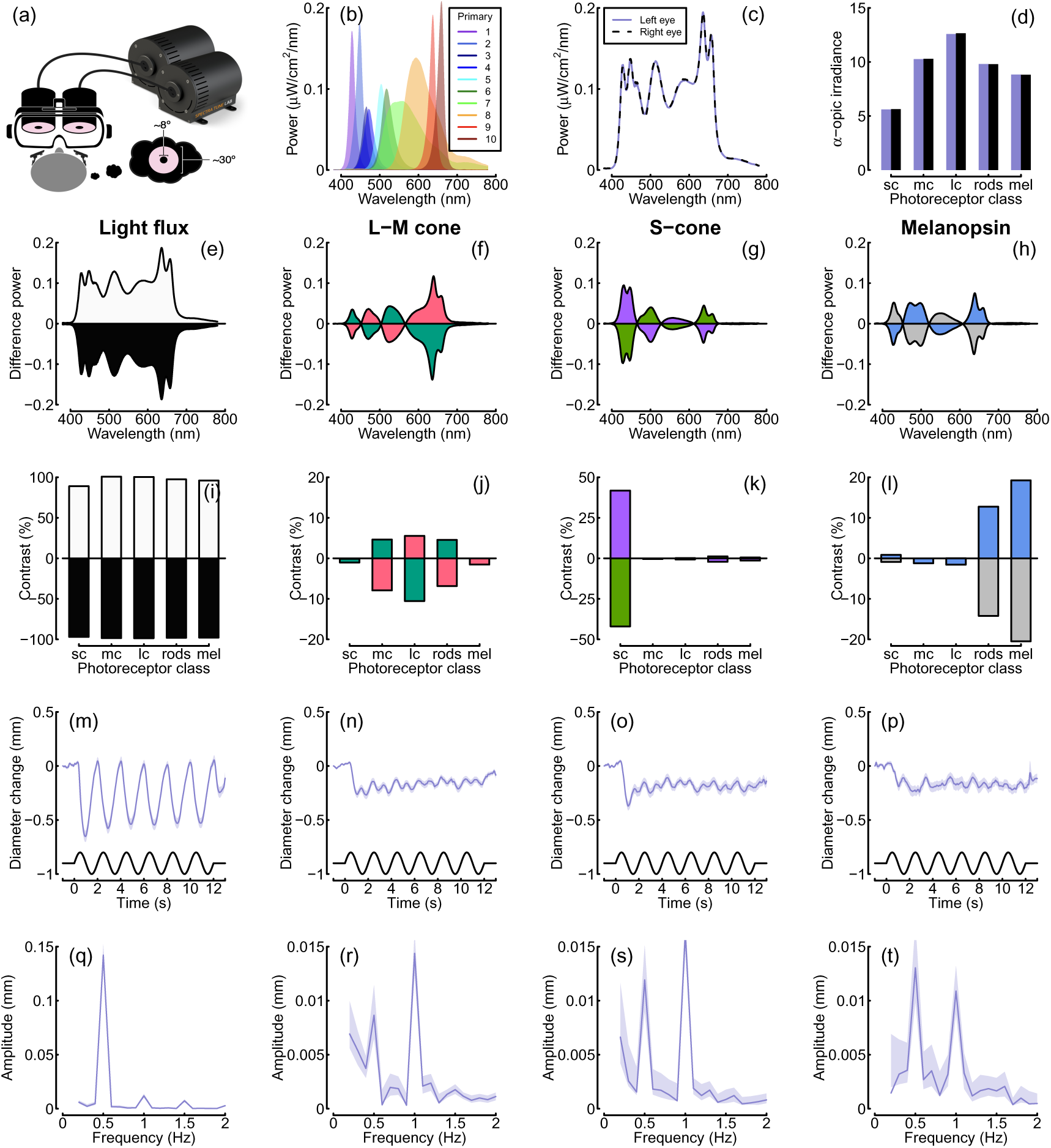
Summary of the spectral power distributions and alpha-opic irradiances for the background and each condition, as well as averaged pupil diameters and Fourier spectra. Panel (a) shows a schematic of the binocular stimulation system for presenting spectrally tuned modulations independently to each eye. The VR headset was attached to a clamp stand that the experimenters could use to adjust the height and align the headset with the eyes of the participant. The participant’s head was supported by a chin rest to keep it in position throughout the experiment. Panel (b) shows the outputs of each LED primary at maximum intensity, and panels (c) and (d) show the overall spectral power distributions and the alpha-opic irradiances of the background spectra used for both eyes. The subsequent rows show the power differences (e-h), and photoreceptor contrasts (i-l) relative to the background, averaged pupil diameter waveforms (m-p) and Fourier spectra (q-t) for binocular stimulation. Column headings indicate the pathway stimulated, and shaded regions in panels m-t indicate bootstrapped 95% confidence intervals.

## Materials and methods

### Participants

Twenty-four participants were recruited for each of the four experiments for a total of ninety-six adult participants (28 male, 68 female), whose ages ranged from 18 to 41. All participants had normal or corrected to normal vision, no known abnormalities of binocular or colour vision, and gave written informed consent. Our procedures were approved by the Ethics Committee of the Department of Psychology at the University of York (identification number 184).

### Apparatus and stimuli

To present synchronised stimulus modulations independently to each eye, two light engines (SpectraTuneLAB, which are thermally stable across time with active cooling according to the manufacturers: LEDMOTIVE Technologies, LLC, Barcelona, Spain), each with 10 independently addressable LED colour channels, were integrated into a customised binocular viewing system. The light engines were operated via a Python interface to their REST API [23], which supports synchronous launch and playback of spectral sequences prepared in advance and stored in JSON format. A special command was commissioned from LEDMOTIVE to allow two different stimulus files to be launched simultaneously from the two devices. The synchronisation of the spectra from the two devices was tested and showed that they were synchronised to within about 3ms.

When preparing the spectral sequences, the age of participants was used to account for the yellowing of the lenses. We used the silent substitution technique [24] to selectively stimulate specific photoreceptor classes. Silent substitution exploits the fact that each photoreceptor class has a distinct spectral tuning that overlaps with the others. Using a multiprimary system, in which the primaries (i.e. LEDs) have different spectra, it is possible to target one class of photoreceptors while maintaining the others at a constant activity level, effectively silencing them [25, 26]. We calculated silent substitution solutions using the PySilSub toolbox [27], using linear algebra. The outputs of the two light engines (see Figure 1b,c) were calibrated using an Ocean Optics Jaz spectroradiometer, which was wavelength-calibrated to an Argon lamp and intensity calibrated using a NIST-traceable light source. Each primary was also linearised using a polynomial fit (see Figure S1 for details). We used the 10-degree cone fundamentals [28], and estimates of melanopsin absorbance spectra from CIE S 026 (discussed in a previous paper [27]) to calculate *α*-opic irradiance.

The output from the light engines was directed through liquid light guides (LLG3-8H: Thorlabs Ltd, Cambridgeshire, UK) and diffused onto semi-opaque and highly diffusive white glass discs with a diameter of 50 mm for even illumination (34-473: Edmund Optics, York, UK). The light guide gaskets were butt-coupled to the light engine diffusers with threaded adapters (SM1A9, AD3LLG: Thorlabs Ltd, Cambridgeshire, UK) and the exiting ends of the light guides were mated with 51 mm depth optical cylinders (SM2L20: Thorlabs Ltd, Cambridgeshire, UK) via appropriately threaded adapters (AD3LLG, SM2A6: Thorlabs Ltd, Cambridgeshire, UK). The stimulus diffuser discs were retained at the front end of the optical cylinders about 51 mm from the light source, at which distance the output beam was sufficiently dispersed to afford even illumination of the diffuser when viewed from the front. To guarantee safe illumination levels, a circular neutral-density filter with the same diameter of the white glass discs (50 mm) and an optical density of 0.6 log units was placed in the optical path between the light source and the diffusers. A small circular piece of blackout material with a diameter of about 8 degrees (10 mm) was positioned centrally on the front of each diffuser disc to aid as a fusion lock, as a fixation point, and to occlude the fovea.

The diffuser discs were positioned in the objective planes of the lenses of a modified VR headset (SHINECON SC-G01, Dongguan Shinecon Industrial Co. Ltd., Guangdong, China), which was used by the participants to view the stimuli. The stimuli were two discs of flickering light with a diameter of about 30 degrees, which were fused together into a cyclopean percept resembling a donut-shaped ring of light, similar to that used in other studies [5–7, 29, 30]. The VR headset modifications allowed for small adjustments to account for individual differences in interpupillary distance and focal length. The use of this set up allowed us to modulate the stimuli in three different ocular configurations, similar to the ones we used in our previous study [3]: monocular, binocular and dichoptic. In the monocular configuration, the unstimulated eye still saw a non-flickering disc of mean light flux. A schematic of the stimulation system is shown in Figure 1a. Pupillometry data were collected using a binocular Pupil Core eye-tracker headset (Pupil Labs GmbH, Berlin, Germany [31]) running at 120 Hz, and the signals were recorded with the Pupil Capture software.

Our previous study [3] used a temporal frequency of 2Hz for foveal luminance flicker, and recorded EEG data simultaneously with pupillometry. Initial pilot experiments indicated that this frequency was too high to elicit measurable responses when stimulating individual photoreceptor pathways. For all experiments, we therefore used a primary flicker frequency of 0.5 Hz, as previous literature showed that this was slow enough to elicit a pupil response from all photoreceptor classes [7]. We also focussed on only recording pupillometry data as this frequency would be too slow to elicit steady-state EEG responses [32].

For all experiments, sinusoidal temporal modulations were presented against the same background spectrum (matched between the eyes), which was used to achieve silent substitution in the three photoreceptor modulation experiments. The background spectra were defined by setting all channels to half maximum output for the brighter of the two devices (STLab 1, left eye) and then using the STLab 1/STLab 2 calibration ratio to find the equivalent settings for the companion device (STLab 2, right eye). The background spectrum illuminance was about 74 lux, or 68.5 cd/m^2^. The spectral power distributions and *α*-opic irradiances of the background spectra for both eyes are shown in Figure 1c-d.

Silent substitution stimuli were prepared and calibrated for each participant with custom Python software [27] and Python scripts. Estimates of photoreceptor spectral sensitivities for each participant were constructed from the known photopigment absorbance spectra [28], taking account of the peak axial density of the respective photopigments, as well as lens [28,33,34] and macular pigment density [34,35], in accordance with the field size and age-dependent CIEPO06 observer model [36]. The melanopic and rhodopic action spectra of the 32-year-old standard observer were taken from CIE S 026 [37] and then adjusted for age-related lens transmittance with a spectral correction function, in line with the standard. Macular pigment correction was not applied to the rhodopic and melanopic action spectra because rods are not present at the fovea and ipRGCs sit above the retinal pigment layer [38].

In the light flux experiment, the stimulus intensity was increased and decreased relative to the background, which we expected to modulate all photoreceptor classes (see Figure 1e,i). In the L-M cone modulation experiment, we used silent substitution to increase the L-cone activity, and simultaneously decrease the M-cone activity, during the first half-cycle of the sine wave. In the second half-cycle the polarity of the modulation reversed (see Figure 1f,j). The maximum available L-M contrast was about 10%. In the S-cone modulation experiment, we increased and decreased S-cone-directed signals, whilst keeping activity in the other photoreceptors constant (see Figure 1g,k). Our system allowed a maximum contrast of 45%. Finally, in the melanopsin experiment, we modulated the activity of the melanopsin-containing intrinsically photoreceptive retinal ganglion cells, whilst keeping cone activity constant (see Figure 1h,l). The maximum available melanopsin contrast was 22%. We assume that the activity of rods was constant at the high background luminance intensity used here, and so did not attempt to silence rod activity in any condition, as this would have greatly reduced the available dynamic range. Splatter on nominally silenced photoreceptors was small (see Figure 1j-l), well below the levels that would be expected to generate measurable pupil modulations, although we can observe that the rods may not be saturated in the L-M and melanopsindirected stimuli (Figure 1j and 1l) and could intrude in these two experiments. We also estimated activation of penumbral L and M cones [5, 29, 39], which was minimal (*≤* 1.5 contrast; less than splatter on the open-field cones) for melanopsin-directed stimuli. We note that the temporal frequency of our modulation (0.5 Hz) is well below the range where penumbral cone activation can elicit visible percepts, and that such percepts fade after around 1 second [29], and do not typically affect pupil responses [7].

### Procedure

Before the start of each experiment, participants adjusted the objective planes of the lenses with the help of the experimenter until the stimulus was in focus and they perceived the two pieces of blackout material as one fused disc. Pupil responses to binocular temporal contrast modulations were examined in a factorial design that combined six ocular conditions and five temporal contrast levels: 6, 12, 24, 48 and 96% of the available dynamic range. This design, similar to that used in our previous studies [3, 40, 41], was applied in four separate experiments, each with a different mode of photoreceptor stimulation. In the first three conditions, the discs flickered at 0.5 Hz, in either a monocular, binocular or dichoptic arrangement. In the dichoptic condition the non-target eye saw a flickering fixed contrast of 48% of the available dynamic range. In the remaining three conditions (the cross-frequency conditions) one eye’s disc flickered at 0.4 Hz, and the other eye’s disc flickered at 0.5 Hz. This included monocular responses at 0.4 Hz, as well as binocular (one eye sees each frequency at the target contrast) and dichoptic (target stimulus flickering at 0.5 Hz, mask contrast of 48% at 0.4 Hz in the other eye) arrangements. We counterbalanced presentation of the target stimulus across the left and right eyes. The full set of conditions is summarised in Table 1.

**Table 1:** The six ocular conditions used in each of the four experiments. The target eye was modulated at the target contrast (one of five levels); the response was measured at the frequency or frequencies indicated. Target-eye assignment was counterbalanced across the left and right eyes.

| Condition | Target eye | Other eye | Analysed at | Purpose |
| --- | --- | --- | --- | --- |
| Monocular (0.5 Hz) | 0.5 Hz, variable | Steady background | 0.5 Hz | Response to one eye |
| Binocular (0.5 Hz) | 0.5 Hz, variable | 0.5 Hz, variable | 0.5 Hz | Binocular facilitation |
| Dichoptic (0.5 Hz) | 0.5 Hz, variable | 0.5 Hz, fixed 48% | 0.5 Hz | Interocular suppression |
| Monocular (0.4 Hz) | 0.4 Hz, variable | Steady background | 0.4 Hz | Frequency control |
| Binocular cross | 0.5 Hz, variable | 0.4 Hz, variable | 0.5 & 0.4 Hz | Cross-frequency interaction |
| Dichoptic cross | 0.5 Hz, variable | 0.4 Hz, fixed 48% | 0.5 & 0.4 Hz | Cross-frequency suppression |

The experiments were conducted in a windowless room, in which the only source of light was the modified VR headset. The participants sat as close as possible to the VR headset, leaving enough space for the eye-tracker to record the eyes. Each experiment was carried out in a single session of around 45-60 minutes, divided into three blocks of 15-17 minutes each. In each block, there were a total of 60 trials lasting 15 seconds each (12s of stimulus presentation, followed by 3s of interstimulus interval). The participants were given no task other than look at the black fixation dot while trying to minimise their blinking during the presentation period. For all experiments other than the light flux condition, participants adapted to the unmodulated background luminance for two minutes before stimulation began.

Before the start of the L-M experiment, participants completed a luminance nulling perceptual calibration procedure in L-M cone space on an Iiyama VisionMaster^TM^ Pro 510 display (800 x 600 pixels, 60 Hz refresh rate). During the task, participants were presented with a disc flickering within the L-M cone space (between magenta and cyan). Using a trackball, participants adjusted the angle in cone space to find their subjective isoluminant point, which resulted in changing the flickering intensity of the stimulus until the amplitude of the flicker appeared to be minimised. The result was used to modify the requested contrasts during stimulus preparation so as to account for individual differences affecting perceived illuminance, principally the L:M cone ratio [42, 43].

### Data analysis

The pupillometry data were analysed using the same method we used in our previous study [3]. The data were converted from mp4 videos to a csv text file using the Pupil Player software [31], which estimated pupil diameter for each eye on each frame using a 3D model of the eyeball. The individual data were then loaded into R for analysis, where a ten-second waveform for each trial in each eye was extracted (excluding the first two seconds after stimulus onset). We interpolated across any dropped or missing frames to ensure regular and continuous sampling over time. The Fourier transform was calculated for each waveform, and all repetitions of each condition were pooled across eye and then averaged. Finally, data were averaged across all participants to obtain the group results. We used coherent averaging and at each stage we excluded data points with a Mahalanobis distance exceeding *D* = 3 from the complex-valued mean [44]. For monocular stimulation, we confirmed that the consensual response was equivalent to the response in the stimulated eye. For all experiments, we used a bootstrapping procedure with 10^4^ iterations to estimate standard errors across participants. All analysis and figure construction was conducted using a single R-script, available online, making this study fully computationally reproducible: https://osf.io/gdvt4/.

### Computational model and parameter estimation

To quantitatively summarise our data, we used the same model described in our previous study [3]. The model has the same general form as the first stage of the contrast gain control model proposed by Meese and colleagues [22], but omits the second stage. For the previous model that we used [3], the exponent of the numerator and denominator had fixed values of 2 and (implicitly) 1. Here, we allow these parameters (called p and q) to be free, to allow different shapes of contrast response function, e.g. accelerating or saturating.

In words, the model works as follows. Each eye produces an excitatory response that grows with the contrast in that eye. This response is then divided by a term that includes the contrast in the other eye, so that stimulating one eye reduces the response to the other. The two suppressed monocular signals are summed to give the binocular response. The weight *ω* sets the strength of this interocular suppression: values below 1 give weak suppression, and therefore stronger binocular facilitation, whereas values near or above 1 give strong suppression, which can produce ocularity invariance or, when suppression is strong enough, a binocular response that is smaller than the monocular one. The responses of the left eye and right eye channels are as follows:

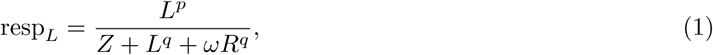

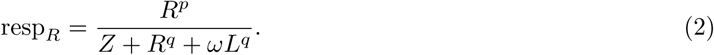

where L and R are the contrast signals from the left and right eyes, p and q are exponents, Z is a saturation constant that shifts the contrast-response function laterally, and *ω* is the weight of suppression from the other eye.

The responses from the two eyes are then summed binocularly:

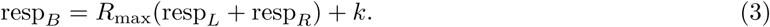

where k is a noise parameter, and Rmax scales the overall response amplitude. The models were fit using a hierarchical Bayesian framework implemented in Stan [45]. The data for each photoreceptor type and response frequency was fit separately, for a total of 8 model fits. We did not fit the two eyes separately. Target-eye assignment was counterbalanced across participants, and responses were pooled across eye before fitting, after confirming that the consensual response matched the response in the stimulated eye (see Data analysis). The prior for the *ω* parameter was Gaussian, with a mean of 1 and standard deviation of 0.5. Priors for the other free parameters were also Gaussian, with mean values based on previous work [3]. All free parameters were constrained to be positive (and the saturation constant *Z* to be at least 1), so these Gaussian priors were truncated at those bounds. The model was hierarchical and was fitted to the individual-participant amplitudes rather than to the group averages: each participant had their own set of parameters, drawn from group-level distributions whose means we report and compare across conditions. We ran 16 chains of 64,500 iterations each, discarded the first 2,000 iterations of each as warm-up, and thinned by a factor of 10, giving 103,200 retained samples per fit (over 10^6^ samples before thinning). The group-level parameters converged well in every fit (split-*R*^^^ *<* 1.02 and effective sample size *>* 3,000). In the melanopsin first-harmonic fit a few participant-level exponent and suppression parameters reached *R*^^^ = 1.06; the group-level suppression weight, which is our parameter of interest, had *R*^^^ = 1.00. As a summary of fit, we report the root-mean-square error between the group-mean data and the model prediction for each fit in Table 2.

**Table 2:** Convergence diagnostics and fit statistics for the eight model fits. Maximum split-*R*^^^ is taken across all parameters; minimum effective sample size (ESS) is taken across the group-level parameters; RMSE is the root-mean-square error between the group-mean data and the model prediction.

| Fit | max $\hat{R}$ | min ESS | RMSE (mm) |
| --- | --- | --- | --- |
| Light flux 1F | 1.00 | 17700 | 0.0043 |
| L-M cone 1F | 1.01 | 9800 | 0.0016 |
| S cone 1F | 1.01 | 6200 | 0.0028 |
| Melanopsin 1F | 1.06 | 10300 | 0.0030 |
| Light flux 2F | 1.01 | 5700 | 0.0014 |
| L-M cone 2F | 1.02 | 3100 | 0.0021 |
| S cone 2F | 1.02 | 3000 | 0.0020 |
| Melanopsin 2F | 1.00 | 11300 | 0.0035 |

## Results

Figure 1m-p shows averaged waveforms of pupil diameter in response to binocular stimulation. For light flux stimuli (Figure 1m) a strong modulation is apparent at the stimulation frequency (0.5 Hz), which is also clear in the Fourier amplitude spectrum (Figure 1q). For binocular stimulation of the L-M pathway, S-cone pathway and melanopsin pathway, pupil modulations were less than 10% of the amplitude of the light flux modulation (note the change in y-axis scale for the Fourier spectra), but still apparent at 0.5 Hz in the Fourier spectra (Figure 1n-p). We also observed a second harmonic response (at 1 Hz) for all conditions, which was weaker than the first harmonic for light flux and melanopsin stimulation, but stronger for L-M and S-cone stimulation. The second harmonic is also apparent in the pupil waveforms shown in Figure 1n-p. Our main analysis therefore focuses on the amplitude of the pupil modulations at both the first and second harmonic frequencies across different stimulus conditions.

A response at twice the stimulus frequency tells us that the pupil does not simply follow the sinusoidal modulation. A purely linear system driven by a sinusoid responds only at the stimulus frequency, so energy at the second harmonic must reflect a nonlinear transformation of the input. Such harmonic distortion can arise from many sources including rectification or asymmetry between constriction and redilation. We therefore treat the second harmonic as a complementary measure of nonlinear temporal processing in the pupil response, rather than as a separate oscillatory mechanism. This analysis was exploratory, as we had no specific prediction about the size of the second harmonic in each condition.

Figure 2a-d shows contrast response functions across stimulation conditions for responses at the first harmonic of the main stimulation frequency (0.5 Hz). In each plot, the response to monocular stimulation is given by the red circles and typically increases monotonically as a function of stimulus (temporal) contrast. Relative to monocular stimulation, binocular stimulation led to higher response amplitudes, indicating a binocular facilitation effect, for the light flux and S-cone conditions (Figure 2a,c), and to some extent for the melanopsin condition (Figure 2d). But the L-M cone condition (Figure 2b) produced a binocular suppression effect, where the response to binocular stimulation was weaker than the response to monocular stimulation (blue squares below red circles). These results indicate that the magnitude of binocular facilitation differs across photoreceptor pathway, suggesting heterogeneity in the underlying neural computation.

**Figure 2:**
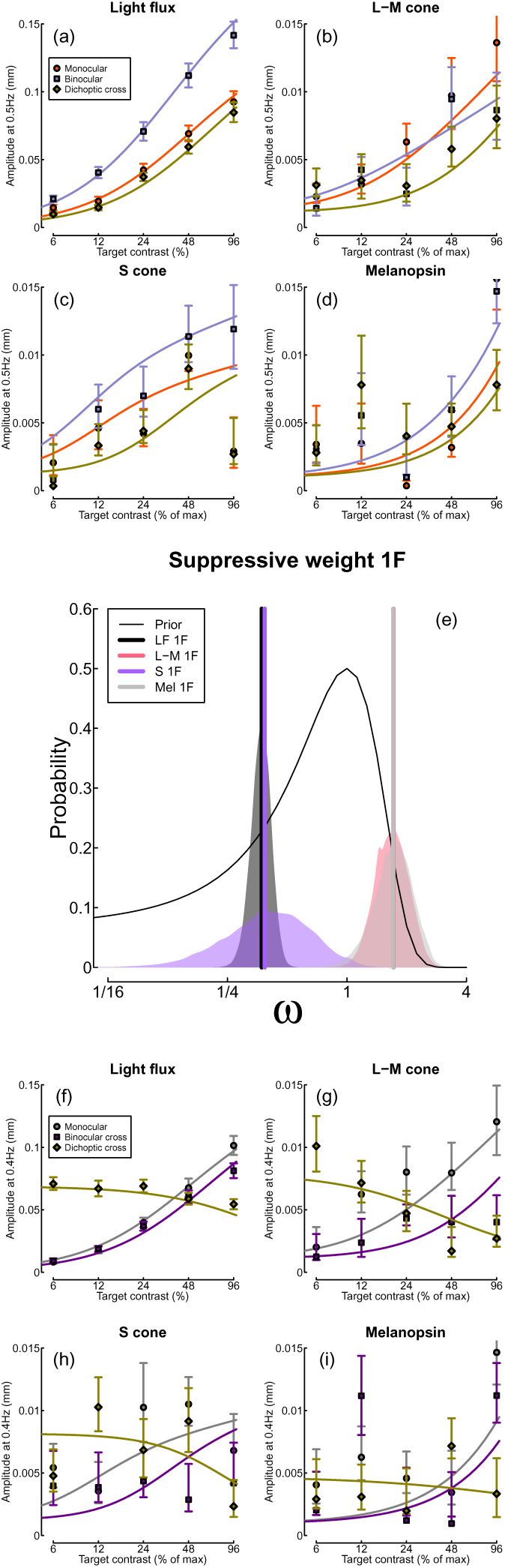
Contrast response functions for pupil modulations in response to flicker at 0.5 Hz (panels a-d), and 0.4 Hz (panels f-i), and posterior parameter distributions for the weight of interocular suppression (panel e). Within each panel (except panel e), data points are the coherently averaged amplitudes for each condition, and error bars indicate bootstrapped 95% confidence intervals. Curves show model fits using the maximum a posteriori (MAP) parameter values (see Table 3). In panel (e), vertical lines show the MAP estimates, and the thin black curve indicates the prior (note the logarithmic x-axis).

**Table 3:** Summary of maximum a posteriori (MAP) parameter estimates for each data set.

| Experiment | $Z$ | $k$ | $\omega$ | $p$ | $q$ | $R_{\max}$ |
| --- | --- | --- | --- | --- | --- | --- |
| Light flux 1F | 95.77 | 0.00083 | 0.37 | 1.32 | 1.24 | 0.0905 |
| L-M cone 1F | 49.82 | 0.00109 | 1.71 | 1.50 | 1.21 | 0.0032 |
| S cone 1F | 63.00 | 0.00121 | 0.39 | 1.94 | 1.79 | 0.0042 |
| Melanopsin 1F | 79.61 | 0.00094 | 1.71 | 1.29 | 0.76 | 0.0025 |
| Light flux 2F | 48.81 | 0.00089 | 1.24 | 1.15 | 0.88 | 0.0039 |
| L-M cone 2F | 66.00 | 0.00075 | 0.95 | 1.10 | 0.82 | 0.0050 |
| S cone 2F | 75.94 | 0.00097 | 2.22 | 1.41 | 0.83 | 0.0018 |
| Melanopsin 2F | 67.94 | 0.00101 | 1.37 | 0.90 | 0.49 | 0.0035 |

In contemporary models of binocular signal combination, the amount of binocular facilitation is determined by the magnitude of interocular suppression, with strong suppression reducing facilitation [22]. We can estimate the strength of interocular suppression by measuring how much monocular responses are reduced when a dichoptic ‘mask’ is shown to the other eye. In our paradigm, the two components flickered at different frequencies (0.5 and 0.4 Hz) so that their responses remained distinct in the Fourier spectrum [46] (all conditions are summarised in Table 1). The yellow diamond symbols in Figure 2a-d show the target responses in this condition, and in most cases were weaker than the monocular responses (red circles). The strongest dichoptic masking is found in the L-M condition, where we also observed the binocular suppression effect. Suppression can also be estimated from the responses at 0.4 Hz (Figure 2f-i). The reduced amplitude in the binocular cross condition (where the two eyes received different temporal frequencies; purple squares) relative to the 0.4 Hz monocular condition (grey circles), and the progressive decline in amplitude of the dichoptic cross response (yellow diamonds) also differ across photoreceptor conditions, showing similar differences to those observed at 0.5 Hz.

To estimate the extent of interocular suppression for each photoreceptor pathway, we fitted each data set using a Bayesian hierarchical implementation of a simple binocular combination model [3, 22]. Our primary objective was to compare posterior distributions of the weight of interocular suppression, which are shown in

Figure 2e. Consistent with our earlier observations, the strongest suppressive weight corresponds to the L-M and Melanopsin conditions, and the weakest suppression corresponds to the light flux and S-cone conditions, with virtually no overlap between the posterior distributions for weak and strong suppression. The model captured the main features of the contrast response functions (Figure 2a-d,f-i), fitting the light flux data most closely and if fitted the noisier S-cone data least well; it did not capture every data point, and some scatter remains in the weaker responses. Fit statistics and convergence diagnostics for all eight fits are given in Table 2, and a posterior predictive check (Figure S3) plots the model’s 95% predictive intervals against the data for every condition: most points fall within these intervals, with the widest intervals and the largest deviations in the noisier, low-amplitude conditions. The fitted model parameters are given in Table 3.

At the second harmonic frequencies, the levels of suppression were more uniform across different photoreceptor pathways (see Figure 3). In general, suppression estimates were near or above 1 (see lower rows of Table 3), with substantial overlap between the posterior distributions (Figure 3e). The contrast response functions also looked more uniform, and generally involved less binocular facilitation and more interocular suppression than were seen at the first harmonics. The L-M condition now featured the weakest suppressive weight, and the strongest binocular facilitation, which was the opposite pattern seen at the first harmonic.

**Figure 3:**
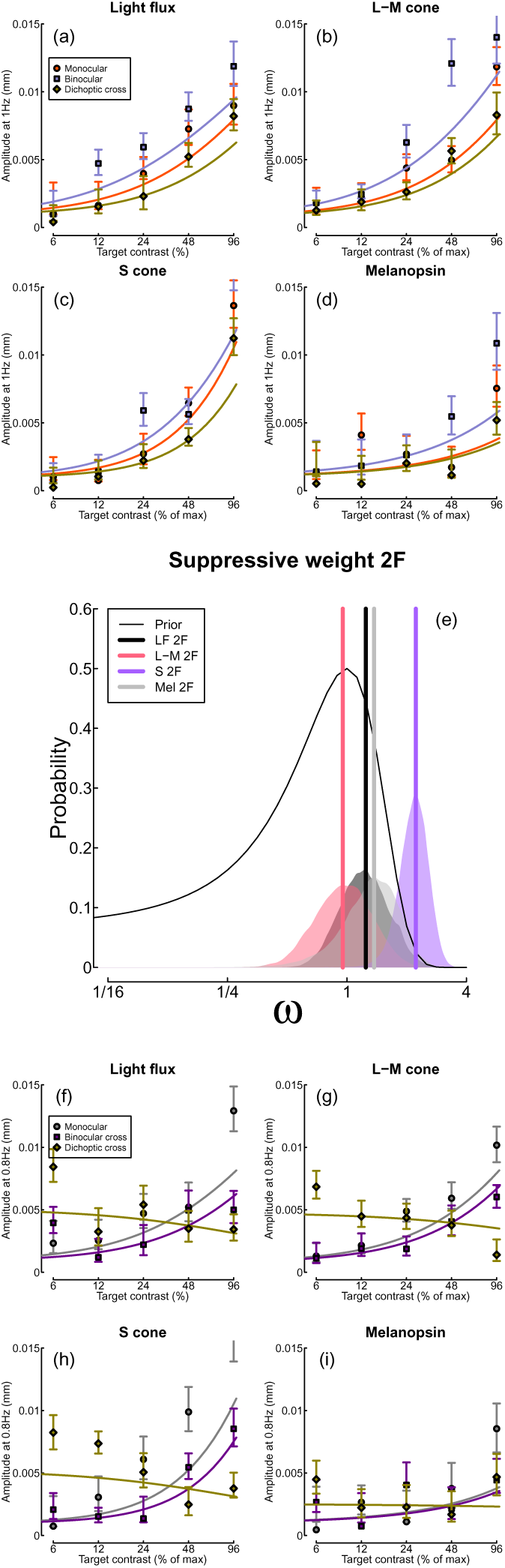
Responses at the second harmonic frequencies (1 Hz and 0.8 Hz), in the same format as Figure 2.

Because response energy can shift between the first and second harmonics depending on the exact shape of the transducer, either harmonic on its own could misrepresent the total periodic response. As a check, we combined the two harmonics into a single measure of periodic response magnitude: the root-sum-square of the first- and second-harmonic amplitudes (Figure S2). The combined measure preserved the main qualitative pattern. Binocular stimulation exceeded monocular stimulation for the light flux and S-cone conditions (binocular facilitation), whereas the L-M and melanopsin conditions showed little facilitation and sat close to ocularity invariance. Combining the harmonics moderated the most extreme ratios, in particular the large S-cone facilitation and the L-M binocular suppression seen at the first harmonic alone, but did not fundamentally alter our conclusions.

Finally, we inspected the response phase of our four stimulation conditions, given previous reports that these differ across pathways [7], and may be in antiphase for melanopsin and S-cone signals. Figure 4 shows the phase angles for the first (a) and second (b) harmonic frequencies for binocular stimulation (monocular stimulation produced very similar results). At the first harmonic, melanopsin and S-cone signals differed in phase by more than 90 degrees, though they were not fully in antiphase. The light flux and L-M responses were approximately in antiphase to each other, but this is likely due to our choice to modulate L+M*−* in the first half-cycle of the sine wave (corresponding to a luminance increase in the light flux condition), and L*−*M+ in the second half-cycle (corresponding to a luminance decrease in the light flux condition). Had we reversed this phase arrangement, we would likely have seen a close phase correspondence between the L-M and light flux conditions (another way to think about this is that L-cone decreases and M-cone increases are processed like luminance increases). Phase differences between the light flux, S-cone and melanopsin conditions are likely to reflect, at least in part, different lags in phototransduction at the earliest stage (i.e. in the retina), although downstream temporal filtering within the pupillary pathway could also contribute. At the second harmonic frequency (Figure 4b), the light flux condition was again out of phase with the other three, and was approximately in quadrature phase with the L-M condition, and in antiphase with both the S-cone and melanopsin conditions (which were in phase with each other). We note that the marked phase differences between conditions make the possibility that our results are dominated by rod activity relatively unlikely (e.g. in the L-M and melanopsin conditions, which have very different phase profiles).

**Figure 4:**
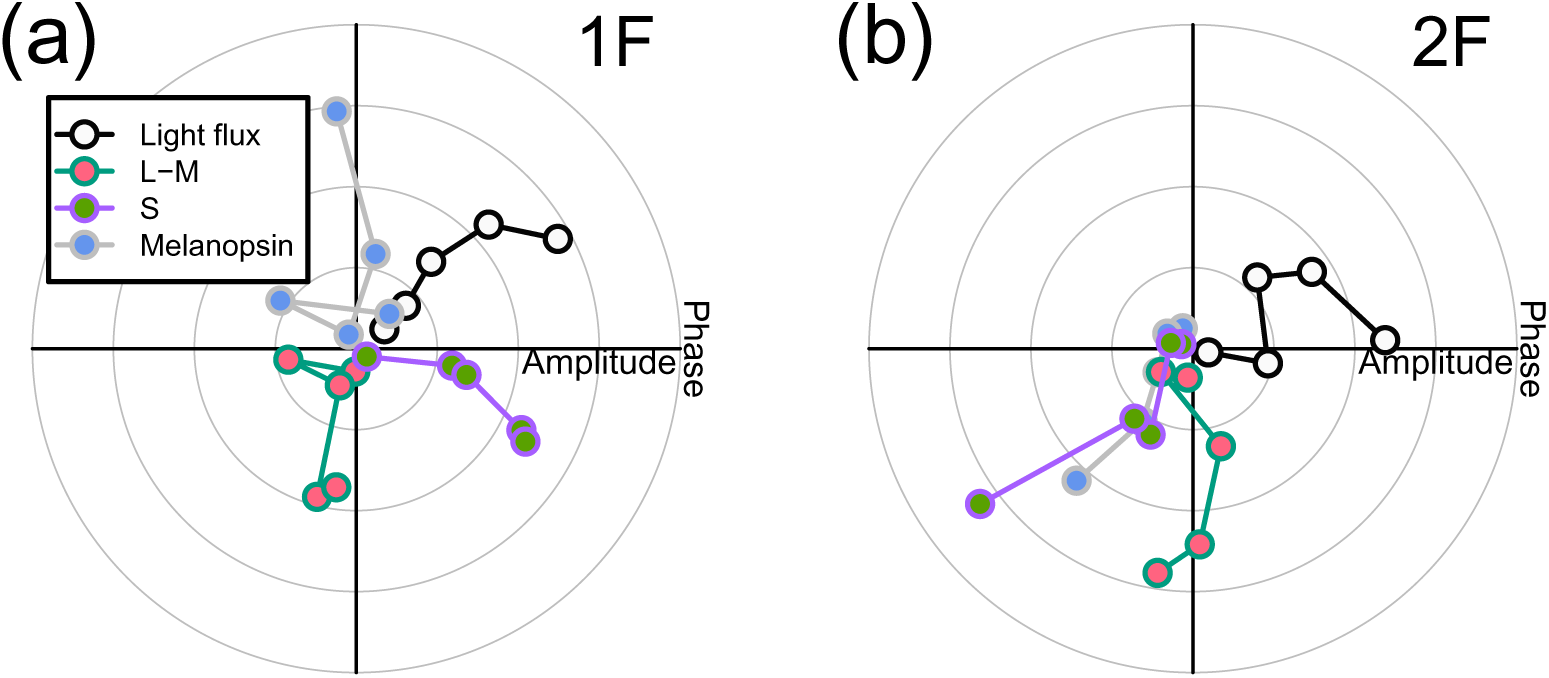
Pupil phase plots at the first and second harmonic frequencies for the light flux, melanopsin, L-M pathway and the S-cone pathway conditions. Panel (a) shows the pupil response at the first harmonic frequency during binocular stimulation. Panel (b) shows the pupil response at the second harmonic frequency during binocular stimulation. The five data points for each pathway correspond to the five contrast levels displayed. In panel (a) the light flux amplitudes have been scaled down by a factor of 10 to enable comparison with the other conditions.

## Discussion

We used binocular pupillometry and silent substitution to measure monocular and binocular responses of the pupils to flickering stimuli when stimulating specific photoreceptor pathways. In all four experiments, we were able to record contrast response functions at both the first and the second harmonic frequencies. All experiments showed that binocular combination in the autonomic nervous system happens in a non-linear manner, with evidence of different magnitudes of interocular suppression depending on the photoreceptor pathway. This pattern of results was confirmed by a computational model, which allowed us to compare the weight of interocular suppression for each pathway. We found that at the first harmonic frequency the L-M and melanopsin pathways involved strong suppression, whereas the light flux and S-cone pathways involved weaker suppression. Suppression was strong in all four pathways at the second harmonic frequency. Finally, the phase of the pupil response revealed different lag times for the different pathways at the first and second harmonic frequencies.

This is the first study to investigate binocular interactions in the melanopsin pathway directly. A previous analysis [47] indicated that there may be a substantial binocular facilitation effect in the circadian pathway, as indexed by melatonin suppression (melatonin is a hormone released by the pineal gland; its production is suppressed by exposure to bright light, particularly when the melanopsin-containing ipRGCs are stimulated). In brief, monocular stimulation of the ipRGCs requires up to ten times the signal strength to produce an equivalent effect to binocular stimulation. This superadditive effect implies an absence of interocular suppression, and perhaps the presence of either a highly compressive nonlinearity, or an AND-style neural operation. Our findings here are different - we do not see substantial binocular facilitation effects in response to melanopsin modulation, and our data indicate that interocular suppression is strong. By measuring the full contrast response function, we can also rule out compressive nonlinearities, because the functions accelerate at both the first and second harmonic frequencies. The explanation for this difference could be due to different anatomical pathways. Binocular combination in the circadian system likely takes place in the suprachiasmatic nucleus, whereas in the pupil constriction circuit the Edinger- Westphal nucleus is the most likely site of binocular integration [9]. Presumably these anatomical differences, and the practical constraints of the two systems, lead to differences in response.

Our recent psychophysical work [48] has looked at binocular interactions in the L-M and S-cone pathways, and compared these to the light flux pathway. Using a contrast discrimination paradigm with spatial modulations of light flux and colour, we found equally strong interocular suppression in all three pathways [48]. This is rather different from our pupillometry results here, which show weaker suppression in the light flux and S-cone pathways than in the L-M pathway (see Figure 2). But the current experiments involve temporal modulations, which are quite distinct from the spatial modulations used in our previous work [48].

Our other recent work [3] has shown that temporal luminance modulations involve much weaker interocular suppression in the cortical response than do spatial luminance modulations [40]. In more recent work, we used steady-state evoked potentials to measure interocular suppression cortically in chromatic and luminance patterns, and found that it depends on both colour and spatial frequency [49]. These different normalisation processes might reflect different priorities for spatial and temporal vision. Spatial vision aims to fuse images to provide binocular single vision, and benefits from ‘ocularity invariance’ [20], in which visual appearance is constant when viewed with one eye or two. Temporal vision is critical for motion perception, which can involve alternative binocular computations, such as calculating velocity differences between the eyes [50]. Of course the present measurements were of pupil size, which may be subject to different anatomical and functional constraints from those in the cortex. Future research should aim to extend our current findings to perception using psychophysical approaches.

### Physiological interpretation of the pathway and harmonic differences

Our four experiments modulate different photoreceptor classes, and the differences between them are open to physiological interpretation. We are cautious about attributing responses to discrete physiological mechamisms because silent substitution isolates photoreceptor contrast rather than an anatomically separate neural pathway: the targeted photoreceptor signals converge with others downstream, and we did not explicitly silence the rods although their contribution at the mean luminance levels used is expected to be small.

The light flux stimulus modulates several photoreceptor classes together and produced by far the largest pupil response. Its weak first-harmonic suppression is consistent with strong binocular facilitation, but because several receptors are co-modulated this condition cannot tell us which retinal input dominates. The isoluminant L-M stimulus targets a cone-opponent signal rather than luminance, and showed the strongest first-harmonic suppression, with a binocular response weaker than the monocular one. This may indicate stronger binocular normalisation for cone-opponent input, though our data cannot localise whether this interaction arises in retinal, pretectal or other subcortical circuitry.

The S-cone stimulus produced a relatively large second harmonic. This could reflect nonlinearity or asymmetry in how S-cone increments and decrements are converted into a pupil response, rather than a separate oscillatory mechanism. For example, it is well known that there are asymmetries in the number of S-cone on- and off- bipolar cells in the retina [51, 52]. The melanopsin-directed responses were delayed in phase relative to the cone-directed responses, which is broadly consistent with the slower phototransduction of the ipRGCs. But the ipRGCs also receive cone and rod input, and our stimulus isolates melanopsin contrast rather than an anatomically independent pathway, so the delay need not arise from melanopsin phototransduction alone.

The meaning of the differences we observe between the 0.4 and 0.5 Hz responses is unclear. The pupil is a temporally filtered and nonlinear system, so a modest change in frequency can alter response phase, waveform asymmetry and harmonic content. Some of the difference may also reflect noisier estimates or imperfect model fit. We therefore do not attribute the 0.4 versus 0.5 Hz differences to distinct physiological mechanisms. A limitation that applies across all of these comparisons is that the four experiments used separate groups of 24 participants, so differences between the photoreceptor-directed conditions are partly between-group differences.

## Conclusions

We have shown that binocular combination of temporal flickering light in the autonomic nervous system depends on which photoreceptors are stimulated. We were able to elicit pupil responses by stimulating the periphery of the retina, and to record contrast response functions for all four photoreceptor-directed conditions. While all conditions showed non-linear combination, they varied in how the signals are combined, particularly in the weight of interocular suppression. This was strong (*ω ≥* 1) for L-M and melanopsin signals at the first harmonic, and all pathways at the second harmonic. Suppression was weaker (*ω* < 1) in the light flux pathway (consistent with previous work), and also for S-cone directed modulations.

## Supporting information

**Figure S1:**
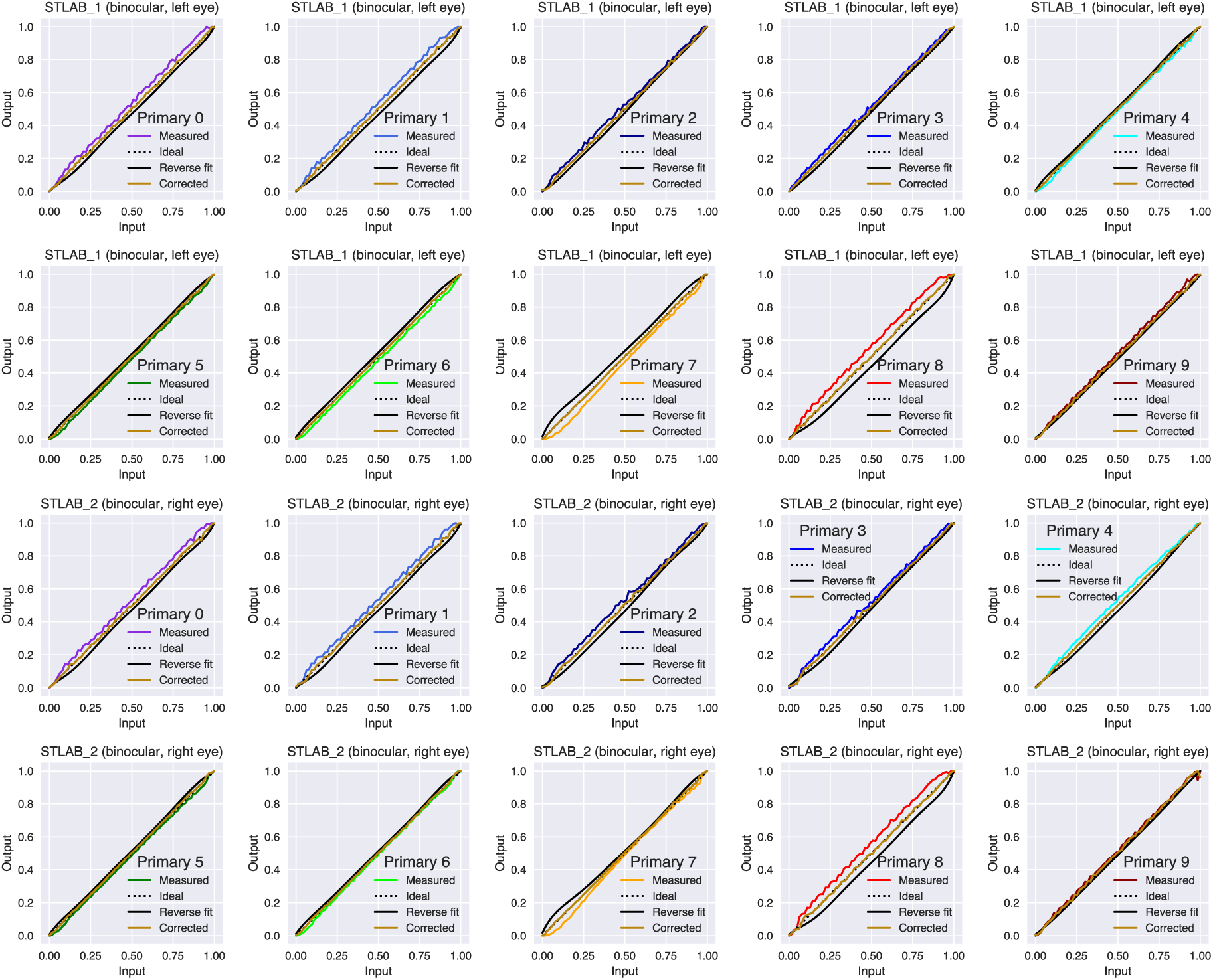
Linearity of primaries for each device. The calibration measures were summed (i.e., total un-weighted irradiance), and the input-output relationship summarised by a 7th order polynomial reverse curve fit. By applying the coefficients of the regression, it is possible to achieve a linear output in the primaries.

**Figure S2:**
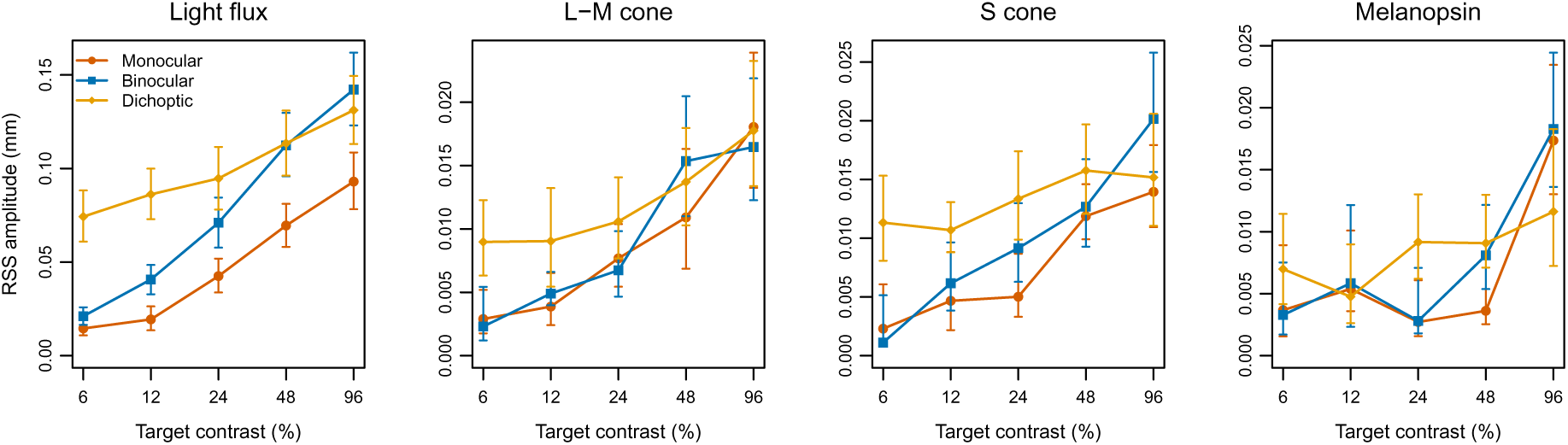
Combined first- and second-harmonic pupil responses. For each photoreceptor-directed condition, the root-sum-square amplitude of the first (0.5 Hz) and second (1 Hz) harmonic is plotted against target contrast, for monocular, binocular and dichoptic stimulation. Points show the coherently averaged group amplitude and error bars show bootstrapped 95% confidence intervals (10^4^ iterations). Combining the harmonics preserves the binocular facilitation seen for light flux and S-cone stimulation, and the near-invariance seen for L-M and melanopsin stimulation.

**Figure S3:**
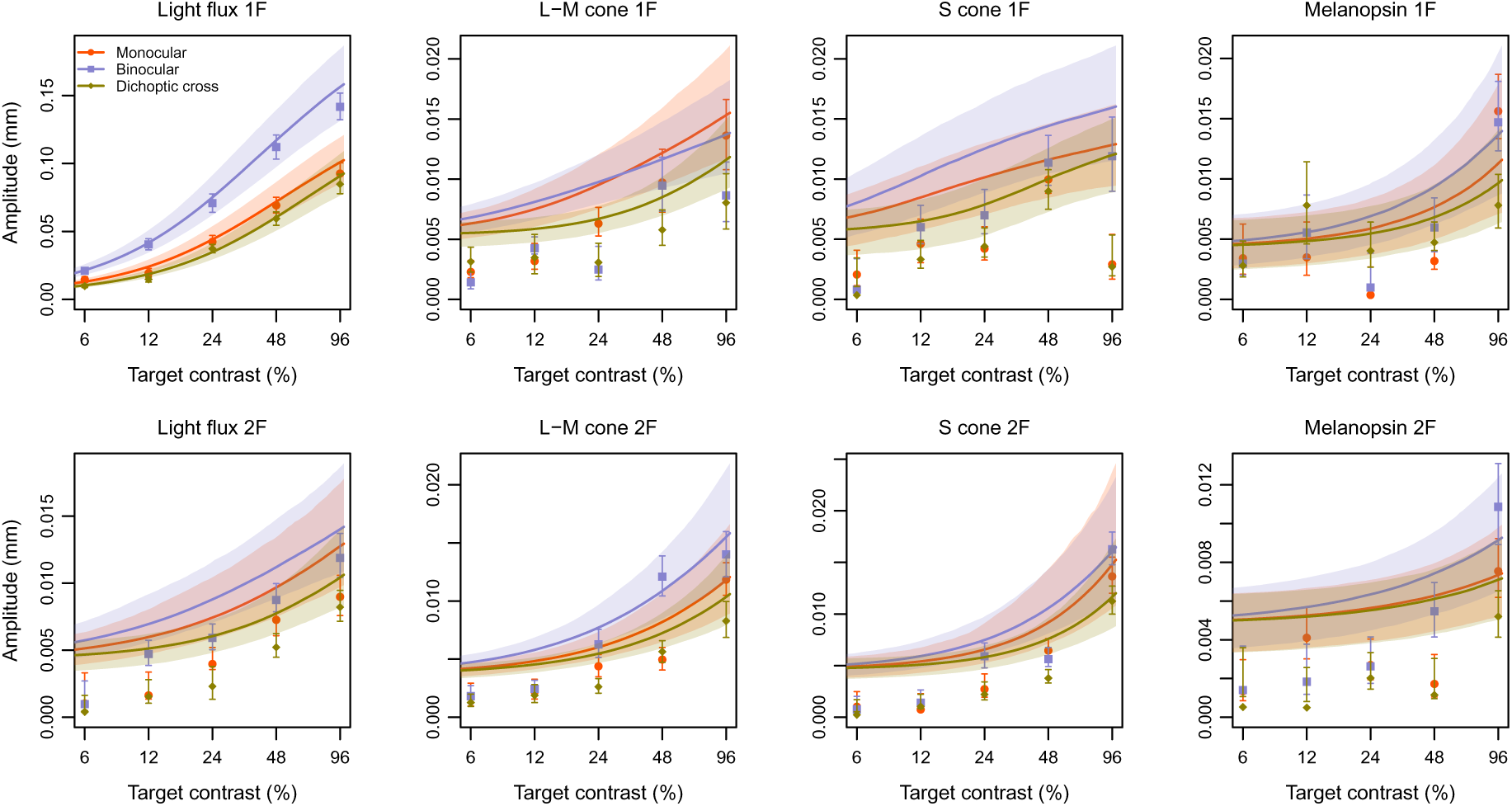
Posterior predictive check for the eight model fits. Each panel plots the observed group-mean amplitudes (points, with bootstrapped 95% confidence intervals) for the monocular, binocular and dichoptic-cross conditions against target contrast, at the first harmonic (top row) and second harmonic (bottom row) for each photoreceptor-directed condition. Solid lines show the posterior predictive median and shaded regions the 95% posterior predictive interval, computed from the group-level parameters. Most data points fall within the predictive intervals; the intervals are widest, and deviations largest, for the noisier low-amplitude conditions.

